# FastTrack, A Strategy to Shorten Time to Degree

**DOI:** 10.1101/2020.06.09.142448

**Authors:** Michael D. Schaller, Mariette Barbier

## Abstract

Recent reports express concern about the sustainability of the biomedical research enterprise in its current form. Recurring concerns include the predictability and sustainability of funding for research, regulatory burden and training the next generation in the biomedical workforce. One specific concern is the duration of training periods during pre-doctoral and post-doctoral studies. This article addresses the issue of time-to-degree (TTD) for doctorates. Many reports stress the importance of shortening the TTD, but provide no recommendations to achieve this goal. Herein, factors potentially affecting TTD are discussed and one mechanism that harmonizes undergraduate and graduate programs is proposed as a strategy to reduce the TTD.

---

The sustainability of the biomedical research workforce has received considerable attention in the last few years, as the government, universities, professional organizations and leaders in the field grapple with establishing guidelines and policies to meet our future needs in a changing research climate [1-4]. Much of the discussion focuses on public research universities since 62% of biomedical research is performed at these institutions, which also train 70% of scientists, engineers, teachers and doctors [5]. One point of concern is the changing demographics of the faculty at research universities and related concern of increased average age of junior investigators to establish their independently funded research programs [6-9]. While multiple factors underlie these changes, recommendations to redress these issues include addressing the long training period required prior to securing a faculty position, and suggest reducing the period of postdoctoral training, but also to shorten the time required for completion of doctoral studies, i.e. the time-to-degree (TTD) [2, 4, 5, 7, 10]. A second point of concern is the change in career paths that current trainees will pursue compared with their predecessors, since the supply of PhDs produced far exceeds the demand for academic research positions [2, 11, 12]. This has led to recommendations for expanding graduate and postdoctoral training to include professional development in preparation for a variety of career outcomes and for increased transparency in training program outcomes to allow program applicants an informed choice [2, 3, 13, 14]. One of the metrics included in all calls for transparency in graduate programs is TTD [2, 3, 9]. The Association of American Universities (AAU) has endorsed calls for transparency, establishing the expectation, but not mandating transparency on doctoral degree outcomes [15]. A group of prominent universities has established the Coalition for the Next Generation of Life Science, which has adopted recommendations for collecting and disseminating program outcomes, including TTD [16]. As this data emerges, programs with shorter TTD will presumably gain an advantage, incentivizing all programs to develop strategies to shorten TTD. The National Academies of Science, Engineering and Medicine recently made recommendations for training STEM graduate students in the 21^st^ century, proposing the necessity of a more student-centric approach to doctoral training and shortening TTD would serve the best interests of the students [3]. Shortening TTD may also increase cost-effectiveness, a mechanism to improve stewardship of public dollars by research institutions [5]. Thus, there are a number of compelling reasons to consider the length of TTD and potential mechanisms to decrease TTD. The challenge is to decrease the time without compromising the quality of the PhD, even as curricula are changing to incorporate additional activities related to career development.

## How long does it take to get a PhD?

The National Science Foundation’s Survey of Earned Doctorates compiles data for TTD for broad areas of study at the national level [17]. The most recent data indicate a median TTD for doctorates in the Life Sciences (all biological sciences and biomedical sciences) of 6.8 years. Over the course of 25 years, this represents a decline of one full year in the time to complete a doctorate (see Figure 1). This data is insufficiently granular to provide insight into specific fields and programs. In 2011, the National Research Council (NRC) published an assessment of the biomedical sciences doctoral programs, compiling data provided by universities, programs, faculty and students and reported on 982 programs across 11 disciplines [18]. The median TTD for 2005-2006 ranged from 5.13 years (Physiology) to 5.68 years (Neuroscience). The Coalition for the Next Generation of Life Science has begun the initiative of dissemination of doctoral program metrics, compiling data at the university and program level at each coalition institution [16]. Using Biochemistry, Microbiology and Immunology programs as examples, the three-year median time to degree since 2010 has ranged from 4.7 to 6.73 years (see Figure 1). The apparent discrepancy between the NRC data, CNGLS data and NSF data could reflect variation in specific programs relative to the median of broadly binned programs, but could also reflect different measures of TTD. The Survey of Earned Doctorates reports the median time to degree since beginning graduate school for graduates in 2017 was 6.8 years and the time since beginning the doctorate program was 5.8 years [17]. While the latter more accurately reflects the doctoral program outcome, this data is not available for previous years precluding longitudinal comparisons. This also illustrates the necessity of standardization of the data collected as variation in the measures can confound interpretation, e.g. time-to-degree vs registered-time-to-degree or defining completion of degree as the dissertation defense vs the date of graduation. Recommendations for increased transparency in outcomes should include guidelines for measures to allow authentic comparisons across institutions.

**Figure 1.**
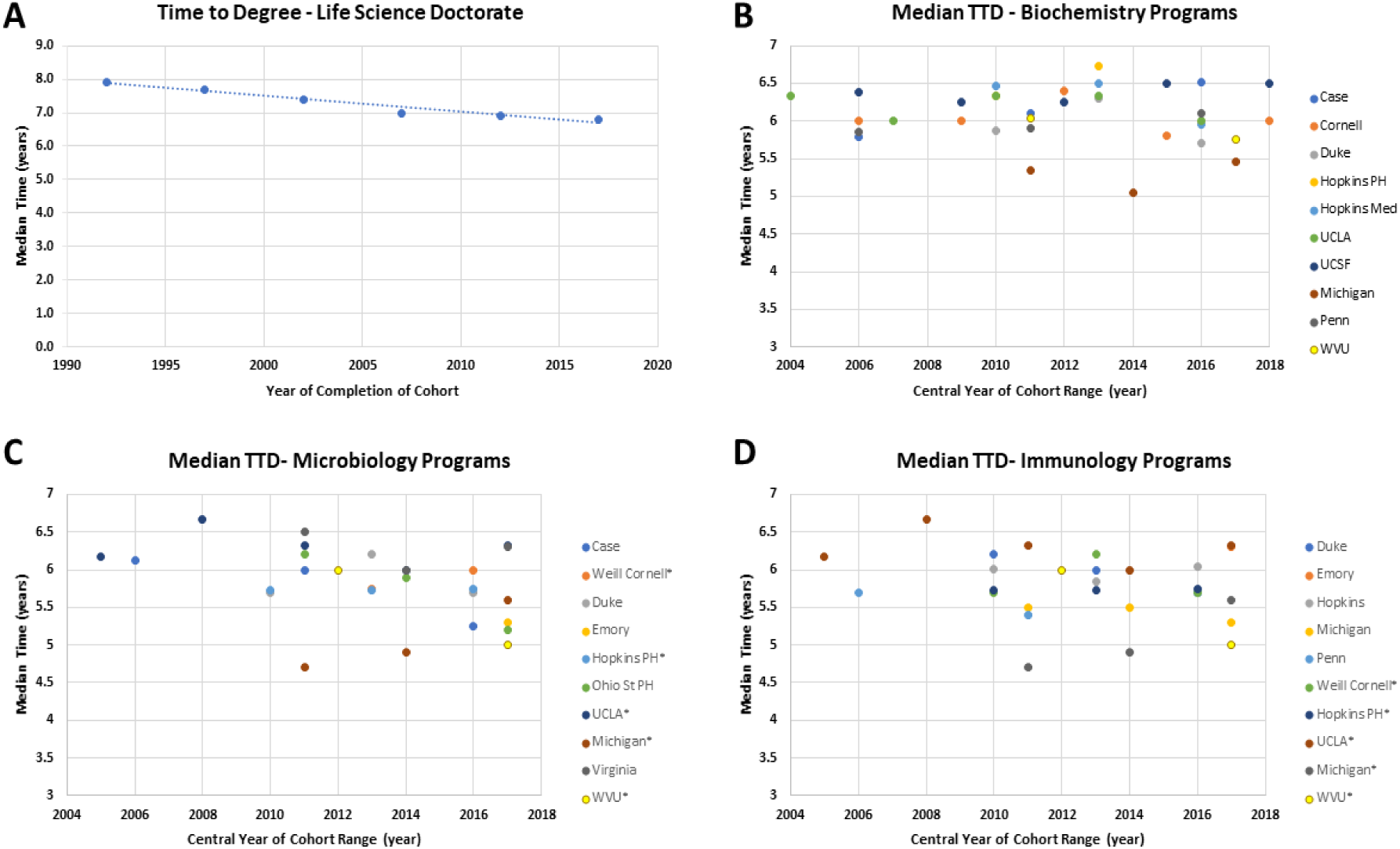
Time to doctorate degree. **A)** Median time to degree for doctorates in the Life Sciences as reported in the National Science Foundation Survey of Earned Doctorates, December 2018. The trend line is plotted. Median time to degree for doctorates in Biochemistry **(B)**, Microbiology **(C)** and Immunology **(D)** as reported by the Coalition for Next Generation Life Science (nglscoalition.org). Data is reported in cohorts of 3 or 5 years. Median is plotted against the middle year of each reported cohort. Median time to degree for doctorates in the BMB and IMP graduate programs at WVU is also shown (two 5-year cohorts are reported). *-indicates combined Microbiology and Immunology Program – data from these programs appears in both C and D.

A linear regression analysis of the median time to degree for Biochemistry, Microbiology and Immunology programs was performed to assess changes in TTD in each field over the last 15 years (see Figure 2). Each curve had a negative slope, but only the slope for the Microbiology programs was significantly different from zero. With the caveat that the data include a small number of programs from a limited number of institutions, there appears to be modest to no change in the median TTD over this period of time. Despite calls for shortening time to degree, there has been little progress over the last 15 years.

**Figure 2.**
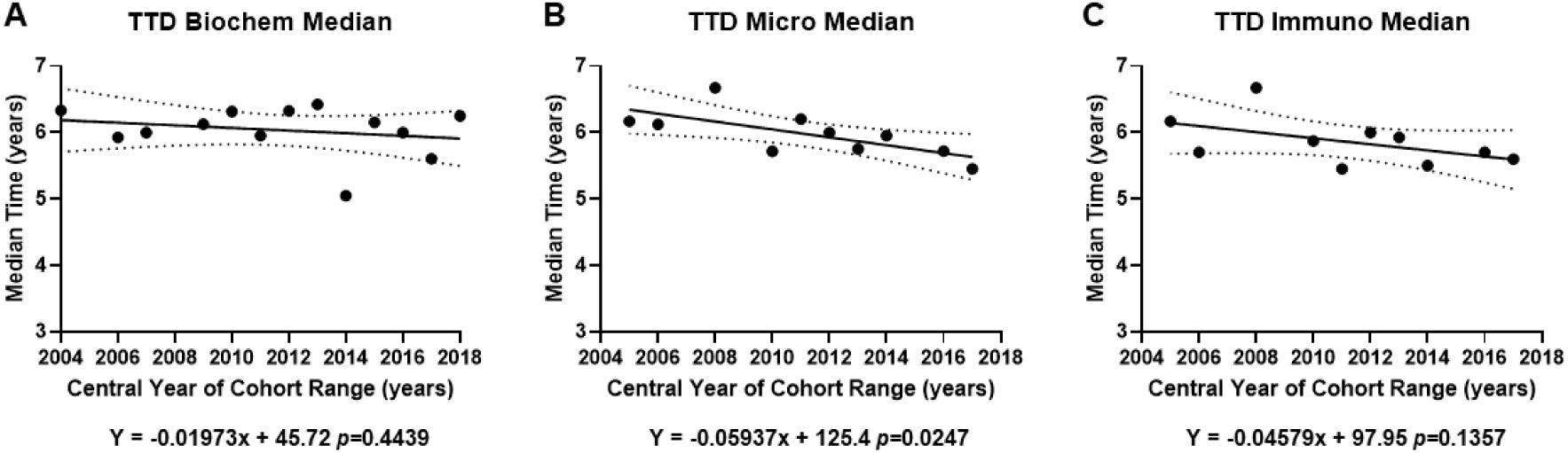
Linear regression analysis of time to doctorate degree. The median time to degree for each year was determined by averaging across all schools for Biochemistry (**A**), Microbiology (**B**) and Immunology (**C**) graduate programs. Similar linear regression lines (solid lines) with 95% confidence intervals (dashed lines) are shown. The equation for each line is also shown. The data was analyzed to determine if the negative slopes were significantly different than zero (p values are shown).

## What Factors are Linked to TTD?

The NRC has published two studies examining factors linked to TTD, the first in 1990 and the second in 2011 [18, 19]. The former study was in response to a rise in TTD observed between the late 1960’s and the 1980’s. For example, the mean total TTD (from receiving a baccalaureate degree) in the biosciences increased from 8.34 +/- 4.76 years to 8.99 +/- 4.06 years between 1967 and 1986. The mean registered TTD, i.e. actual time of study, in the biosciences increased from 5.83 +/- 2.03 years to 6.77 +/- 2.03 years over the same period. The study examined a number of internal and external factors including student attributes, financial support, demographics, market for employment, etc. While individual factors correlated with total TTD, fewer correlated with the registered TTD, and no single factor driving the increased TTD across all programs was identified [19].

The second NRC study examined a number of factors to determine their impact on TTD, attrition rates and the diversity of doctoral programs in 11 disciplines in the biomedical sciences [18]. These factors were primarily internal and included faculty productivity, faculty funding, program size and sources of student funding. There were no compelling correlations drawn between these factors and TTD. Interestingly, there was a correlation between productivity and a **longer** TTD in 6 of the disciplines analyzed [18].

These two studies failed to identify a major factor affecting the TTD in doctoral programs and therefore provided no guidance into strategies to shorten the TTD.

### Recommendations and Strategies to Reduce TTD

Several sources recommend reducing financial support to 5 years providing a financial incentive to shorten the TTD [2, 4]. While the earlier NRC report describes financial support as net neutral [19], other reports suggest financial support has a significant impact on TTD and attrition [20-22], but the outcomes of such a policy are difficult to predict. While a defined period of funding can be a motivation for timely completion, the TTD and attrition rate for students whose studies extend beyond the funding period may increase. Additional structural changes to doctoral programs are likely required to reduce TTD significantly. In their analysis of graduate education in the 21^st^ century, the national academies suggest reviewing all aspects of the curriculum to find ways to shorten TTD, but offers no concrete suggestions [3]. Others have suggested that programs experiment to define best practices.

Mentoring is a key factor in pre-doctoral education [21, 22], and workforce recommendations include suggestions to improve training in mentorship and better engaging dissertation committees in advising and monitoring student progress toward degree completion [22]. While mentorship can clearly impact progression to degree, the mentoring relationship is a partnership between mentor and student, and the students should be engaged and enlightened about mentoring issues. Instruction in ‘mentoring up’ and project management will provide insight into approaching the mentoring relationship from the student’s perspective and provide management skills that may contribute to shortening TTD [23, 24].

Graduate student training in the 21^st^ century has evolved to include training opportunities to prepare trainees for diverse careers, as well as rigor, reproducibility, and responsible conduct of research. Mandated in part by funding agencies, these essential courses and training opportunities add to the time spent in the classroom, creating the need to restructure curricula to integrate these elements without increasing TTD. The Johns Hopkins Bloomberg School of Public Health has initiated a pilot program called the R^3^ program to stress the importance of rigor, responsibility and reproducibility in science [25]. In this restructured curriculum, the required didactic components are courses on critical thinking, scientific reasoning, communication and ethics, and the discipline specific knowledge base is built through independent study and elective coursework. The premise is establishing a strong foundation in the skillset required for performing science in a discipline-independent fashion will increase rigor and reproducibility in the work of the students in the program. The curriculum will also provide training in transferable skills, e.g. communication, which meets the recommendations for better preparation of students for a broad range of careers [2, 3, 13, 26-28]. An additional anticipated benefit is shortening the time to degree by better preparing students to intellectually engage in research prior to initiation of their dissertation research. Given the recent implementation of the program, the projected outcomes have yet to be confirmed, but the R^3^ program is an innovative approach in response to recommendations to develop the biomedical workforce for the future.

## Accelerated Graduate Programs – a Path to Shorten TTD

Shortening the time to degree may require modification of the curriculum to reduce didactic content and/or increasing the productivity of doctoral candidates to shorten the time required to complete the dissertation research project. The challenge is compressing the time of study without compromising the training experience and quality. One strategy to achieve this end is admission of students with a stronger knowledge base and more direct experience in the area of planned dissertation research. The best solution might be coordinated change in both undergraduate and graduate education to streamline progression to the doctoral degree. For practical purposes, this can be achieved most efficiently where the undergraduate and graduate programs are at the same institution. This solution is consistent with a specific recommendation by the National Research Council to restructure doctoral programs to enhance pathways for talented undergraduates [5].

Our graduate council at West Virginia University (WVU) recently approved accelerated tracks in the Biochemistry and Molecular Biology (BMB) Graduate Program and the Immunology and Microbial Pathogenesis (IMP) Graduate Program to pilot these ‘FastTrack’ programs. These programs will admit students with exceptional foundational knowledge in the respective disciplines, eliminating the necessity for introductory graduate courses. Matriculants must also have extensive laboratory experience with their proposed research mentor, eliminating the need for laboratory rotations and accelerating their progress in the lab. These modifications will allow students to meet milestones faster and more rapidly develop their dissertation projects in the lab.

Both the BMB and IMP graduate programs are in the School of Medicine and will engage undergraduate programs at WVU to prepare interested students for FastTrack. The BMB FastTrack will target students in the American Society for Biochemistry and Molecular Biology (ASBMB) accredited undergraduate biochemistry major, which is administered in the WVU Davis College of Agriculture, Natural Resources and Design, while IMP FastTrack will recruit students from the Immunology and Medical Microbiology (IMMB) undergraduate program in the School of Medicine. Eligible students will take advanced undergraduate coursework to lay the foundation of knowledge for the FastTrack graduate programs. For BMB FastTrack, students will take advanced courses in metabolism, protein biochemistry, molecular biology, cell biology and signal transduction. Students intending to pursue the IMP FastTrack will take advanced courses in immunology, microbiology, and vaccinology. Importantly, the required coursework is already part of the undergraduate curricula. Matriculants will use online materials to supplement their knowledge base. To develop the research skills and knowledge required to accelerate through their research project, the students must perform 3 semesters of research in the lab of the intended dissertation supervisor. This will provide the theoretical background and practical experience to facilitate the rapid development of a dissertation project upon matriculation. Students that perform undergraduate research on a topic closely related to their future dissertation research can use this experience to jumpstart their graduate work. Additional skills required for success in graduate school are provided in a writing course, offered by the Davis College, and a journal club, offered by School of Medicine faculty, for students in the undergraduate biochemistry program. Similar courses prepare Immunology and Medical Microbiology majors for IMP FastTrack.

Both FastTrack programs will encourage students to take their preliminary and qualifying examinations early in their tenue in the programs. In the fall semester of their first year, BMB FastTrack students will write their qualifying exams concurrently with students in the regular track, who will be entering their second year in the program (see Table 1). Successful completion of the qualifying exam will advance the BMB FastTrack students into the second-year curriculum, and advanced coursework, in the program. The following summer, the BMB FastTrack students will take the Scientific Writing course, which includes an assignment to write a mock F31 application. In the fall semester, the students will write and defend their dissertation proposal, which is in the format of an F31. The proposal exam will be written concurrently with students in the regular track who are early in the third year in the doctoral program. Students in the IMP FastTrack program will initiate advanced coursework in the Fall and Spring of their first year. They will also complete a grant-focused scientific writing course in the spring or summer of their first year. Students will take a combined preliminary and qualifying exam, which includes a written and an oral examination on advanced knowledge in microbiology and immunology, and the defense of their dissertation proposal, also in the format of an F31 application. This timing will allow the development of a well-designed and vetted F31 application for students in both programs, with the input of the supervisory and dissertation committee for guidance in grantsmanship in time for the NIH December submission deadline. The development of this skillset is aligned with a critical milestone in doctoral studies, which is a recommended strategy to improve the experience. The advancement of key milestones toward completion of doctoral studies by a full year in the BMB and IMP Fasttrack programs is expected to reduce the time required for these students to advance to the next stage in their career (see Table 1).

**Table 1.** Comparison of BMB/IMP Standard and FastTrack Graduate Programs.

|  |  |  |  |
| --- | --- | --- | --- |
| BMB/IMP Standard Track |  | BMB/IMP FastTrack |  |
| 1st Year Fall Semester |  | 1st Year Fall Semester |  |
|  |  | August – Enter BMB or IMP Graduate Program |  |
| August – Enter Undifferentiated Curriculum |  | August – Written Qualifying Exam (BMB) |  |
| BMS747 | Foundations Cont. Biomed. Res. I | BIOC785/MICB785 | Journal Club |
| BMS777 | Foundations Cont. Biomed. Res. II | BIOC702*/MICB782*/MICB791B* | An advanced course |
| BMS706 | Cellular Methods | BMS700 | Scientific Integrity |
| BMS700 | Scientific Integrity | BIOC797/MICB797 | Research |
| BMS702 | Laboratory Rotations |  |  |
| 1st Year Spring Semester |  | 1st Year Spring Semester |  |
| January – Enter BMB or IMP Graduate Program |  |  |  |
| BIOC785/MICB785 | Journal Club | BIOC785/MICB785 | Journal Club |
| BMS715/MICB720* | Molecular Biology/an advanced course | BIOC750*/MICB781* | An advanced course |
| BIOC750*/MICB784B* | An advanced course | BMS701 | Scientific Integrity |
| BMS701 | Scientific Integrity | BIOC797/MICB797 | Research |
| BIOC797/MICB797 | Research | MICB790 | Teaching Practicum |
| 1st Year Summer Semester |  | 1st Year Summer Semester |  |
| BIOC797/MICB797 | Research | BIOC797/MICB797 | Research |
|  |  | BMS720 | Scientific Writing |
| 2nd Year Fall Semester |  | 2nd Year Fall Semester |  |
| Written Qualifying Exam (BMB) |  | Dissertation Proposal (BMB) or Combined Qualifying/ Candidacy exams (IMP) – Admission to Candidacy |  |
| BIOC785/MICB785 | Journal Club | BIOC785/MICB785 | Journal Club |
| BIOC702*/MICB782*/MICB791B* | An advanced course | BIOC797/MICB797 | Research |
| BIOC797/MICB797 | Research | MICB790 | Teaching Practicum |
| MICB790 | Teaching Practicum |  | Additional coursework** |
|  |  | Submit F31 Application to NIH |  |
| 2nd Year Spring Semester |  | Additional Semesters – Fall and Spring |  |
| BIOC785/MICB785 | Journal Club | BIOC785/MICB785 | Journal Club |
| BIOC797/MICB797 | Research | BIOC797/MICB785 | Research |
| MICB781* | An advanced course | BMS707 | Experiential Learning |
| MICB790 | Teaching Practicum |  | Additional coursework** |
|  | Additional coursework** |  |  |
| 2nd Year Summer Semester |  | Additional Semesters - Summer |  |
| BIOC797/MICB785 | Research | BIOC797/MICB797 | Research |
| BMS720 | Scientific Writing |  |  |
|  |  | Complete Research, Write and Defend Dissertation |  |
| 3rd Year Fall Semester |  | * - denotes advanced graduate course. Other advanced courses can be substituted.<br>** - additional coursework is based upon interest of the student or advice of supervisory committee |  |
| Dissertation Proposal (BMB) or Combined Qualifying/ Candidacy exams (IMP) – Admission to Candidacy |  |  |  |
| BIOC785/MICB785 | Journal Club |  |  |
| BIOC797/MICB797 | Research |  |  |
|  | Additional coursework** |  |  |
| Submit F31 Application to NIH |  |  |  |
| Additional Semesters – Fall and Spring |  |  |  |
| BIOC785/MICB785 | Journal Club |  |  |
| BIOC797/MICB797 | Research |  |  |
| BMS707 | Experiential Learning |  |  |
|  | Additional coursework** |  |  |
| Additional Semesters - Summer |  |  |  |
| BIOC797/MICB797 | Research |  |  |
| Complete Research, Write and Defend Dissertation |  |  |  |

## Broader Implementation of Accelerated Programs

While the accelerated program is envisioned to reduce TTD relative to students in the regular program, broader implementation will be required to significantly impact the overall TTD for the graduate program. While the BMB FastTrack program is designed to draw students from the ASBMB accredited undergraduate program at WVU, it could be expanded to draw upon students from other ASBMB accredited biochemistry programs, e.g the biochemistry program at Shepherd University. The research experience requirement could be met through summer internships and a semester long exchange program. Programs such as the West Virginia INBRE program provides summer research experiences at WVU in the department of Biochemistry and Microbiology, Immunology, and Cell Biology to students from smaller colleges in the state. This type of program could provide undergraduate-level research experience in a laboratory that the students plan on joining for graduate research studies. An additional strategy is engagement of colleges with MS programs in Biochemistry, Immunology or Microbiology, since their curricula will also provide a stronger knowledge base to rapidly advance in a doctoral program. MS students will also have additional research experience, albeit not directly related to their planned doctoral work. Alignment of MS and doctoral programs between institutions may provide the impetus for collaborative research and the opportunity for exchange programs between labs providing directly relevant lab experience to the MS student and stimulation of research activity in labs at the MS institution.

## Summary

The emerging consensus regarding the future of our biomedical research workforce supports changes in training, including shortening the TTD. Accomplishing these goals requires re-imagining STEM education and mechanisms streamlining the transition into modified doctoral programs could facilitate these outcomes. Development of collaborative efforts between institutions to streamline graduate education will foster stronger ties between the institutions providing opportunities for further innovation in undergraduate and graduate education. As such, the development of similar programs might be supported by initiatives like the NIGMS Innovative Programs to Enhance Research Training, which has the express goal of creating a highly skilled and diverse biomedical workforce.

## Acknowledgments

Thanks to the members of the Biochemistry and Molecular Biology Graduate Studies Committee, Brad Hillgartner, Roberta Leonardi, David Smith and Peter Stoilov, for critical discussion during development of the BMB FastTrack program. The authors would also like to thank Drs. John Barnett, F. Heath Damron, Cory Robinson, Slawomir Lukomski, and Gordon Meares for their insights during the development of the IMP FastTrack program. MS is director of the Cell & Molecular Biology and Biomedical Engineering Training Program (T32 GM133369). MB is the director of the Immunology and Microbial Pathogenesis graduate program.

